# Large-scale genomic analysis of SARS-CoV-2 Omicron BA.5 emergence in the United States

**DOI:** 10.1101/2024.06.20.599933

**Authors:** Kien Pham, Chrispin Chaguza, Rafael Lopes, Ted Cohen, Emma Taylor-Salmon, Melanie Wilkinson, Volha Katebi, Nathan D. Grubaugh, Verity Hill

## Abstract

The COVID-19 pandemic is marked by the continuing emergence of novel SARS-CoV-2 variants. Questions remain about the mechanisms with which these lineages establish themselves in new geographical areas. In this study, we performed a discrete phylogeographic analysis on ∼19,000 SARS-CoV-2 sequences of Omicron BA.5 lineages between February and June 2022 to better understand how it emerged in different regions of the United States (U.S.). We found that the earliest introductions came from Africa, the putative origin of the variant, but the majority were from Europe, correlating with the high volume of air travelers. Additionally, the analysis revealed extensive domestic transmission between different regions of the U.S. driven by population size and cross-country transmission. Results suggest that most of the within-U.S. spread was between three regions that include California, New York, and Florida. Our results form a framework for understanding novel SARS-CoV-2 variant emergence in the U.S.

## Introduction

As the COVID-19 pandemic enters its fourth year, its causative virus, SARS-CoV-2 has demonstrated the ability to evolve into novel variants. The Omicron (B.1.1.529) variant, detected in late 2021 in southern Africa, was deemed a variant of concern (VOC) by the World Health Organization (WHO) and soon became dominant in the U.S. and the rest of the world (1). Omicron is defined by ∼60 mutations, including 32 in the spike protein, which granted it an evolutionary advantage over co-circulating variants due to enhanced intrinsic transmissibility and immune escape (1–4). New sublineages of Omicron have subsequently emerged, as well as recombinants (5,6). Further, the complex mosaic of immunity in the human population, due to different levels of vaccination and/or prior infection means that the landscape for the emergence of SARS-CoV-2 variants has changed since the start of the pandemic. With ongoing variant emergence, changing patterns of global and domestic spread are vital to understand, as they have significant implications in prevention and mitigation plans.

Recent advances in viral sequencing and phylogenetics enabled the timely use of large-scale phylogenetics to understand SARS-CoV-2 dynamics (7). Studies have been conducted globally, for example in Brazil (8), The Gambia (9), and New Zealand (10), to explore the origins, emergence, and dynamics of SARS-CoV-2 variants throughout the pandemic. In the UK in particular, there have been multiple analyses of national-level spread from major population centers, showing early spread from the origin(s) of introduction, and the seeding and subsequent local transmission in new locations (11–14). Further, in the U.S., previous studies have not only shown the increased risk of importation from other states compared to international origin (15) and the importance of superspreading events at facilitating early transmission (16), but have also explored the impact of international introductions of the Alpha variant (17).

Here, we employ a Bayesian discrete phylogeographic framework to understand the introduction and spread of a novel SARS-CoV-2 lineage into different regions of the U.S. We use Omicron sublineage BA.5, first sampled in the U.S. in early 2022, as the focus for analysis due to its rapid national spread, persistence from mid-2022 to early-2023, and public health importance (**Figure 1**). BA.5 established itself during times of relatively lower SARS-CoV-2 incidence and remained prominent in frequency until the end of 2022 (**Figure 1A**, (18). However, BA.5 never achieved complete dominance in the U.S., and instead co-circulated with other major lineages such as BA.2.12.1, BA.4, and XBB.1 (5). Moreover, BA.5 dissemination occurred on the background of a highly immune population due to vaccination and previous infections with other Omicron sub-lineages (5,19). Newer variants are likely to be introduced onto a similar immune landscape, and so the dynamics of BA.5 introductions and dissemination offer a useful case study for how new lineages may spread across the U.S. Further, as most social and travel restrictions are lifted and data streams become more limited, understanding within-country spread will allow for targeted surveillance activities in the future.

**Figure 1:**
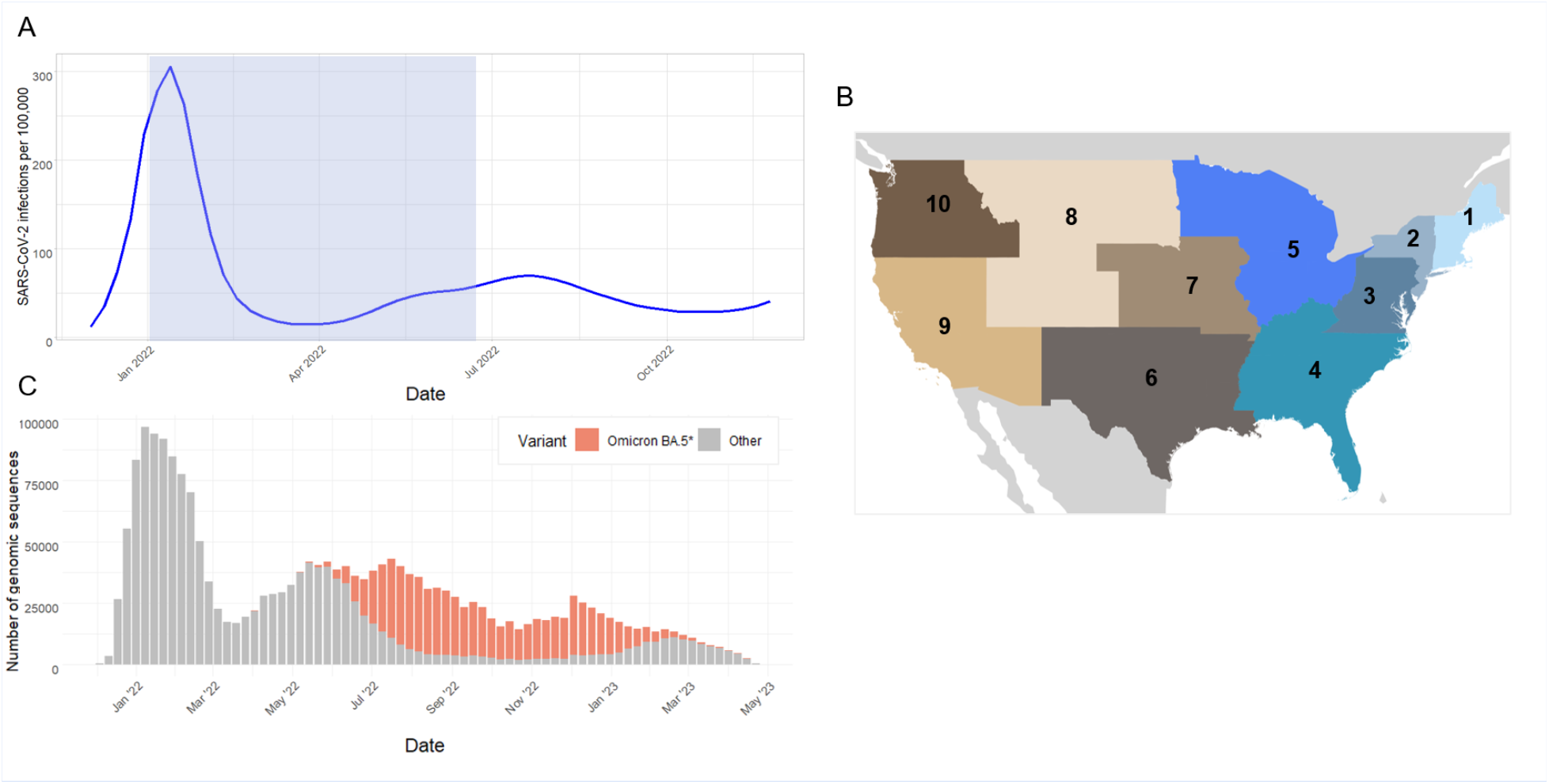
Background information (A) SARS-CoV-2 estimated weekly infections in the U.S. from December 01, 2021, to December 01, 2022, Source: https://www.medrxiv.org/content/10.1101/2020.06.17.20133983v2 (B) Ten regions of the U.S., according to the U.S. Department of Health and Human Services (HHS) designation. (https://www.hhs.gov/about/agencies/regional-offices/index.html) (C) SARS-CoV-2 variant frequency from January 01, 2022, to May 31, 2023.

## Results

### The earliest and largest international introductions of BA.5 Omicron sublineage into the US were from Africa and later ones were from Europe

We began by exploring the dynamics of BA.5 global introductions into the U.S. We performed a discrete phylogeographic analysis at the continent level and between regions within the U.S. (**Figure 1B**) and reconstructed introductions across the resultant phylogeny (**Figure 2A**). We found an average of 1168 (95% CI: 1137-1198) introductions (see **Methods**) from other continents into the U.S. across the whole posterior distribution of the entire period. The inferred time of the first introduction into the U.S. was the second week of February 2022, nearly three weeks prior to the collection date of the earliest U.S. sequence on February 26, 2022 (**Figure 2A, 1C)**. International importation made up most early introductions (68%) before May 2022 (shaded area in **Figure 2B**). This was the case even when, within BA.5’s emergence period, air travel in the U.S. was predominantly between U.S. regions, with domestic volume comprising approximately 80% of all air travel volume (**Figure 4C**). After BA.5 became established by mid-May 2022, 72% of the between-region introductions arose from other U.S. regions (ie. domestic sources; **Figure 2C**). Many international introductions came from Asia (27.8%), Europe (26.3%), and Africa (14.7%) **(Figure 3A**).

**Figure 2:**
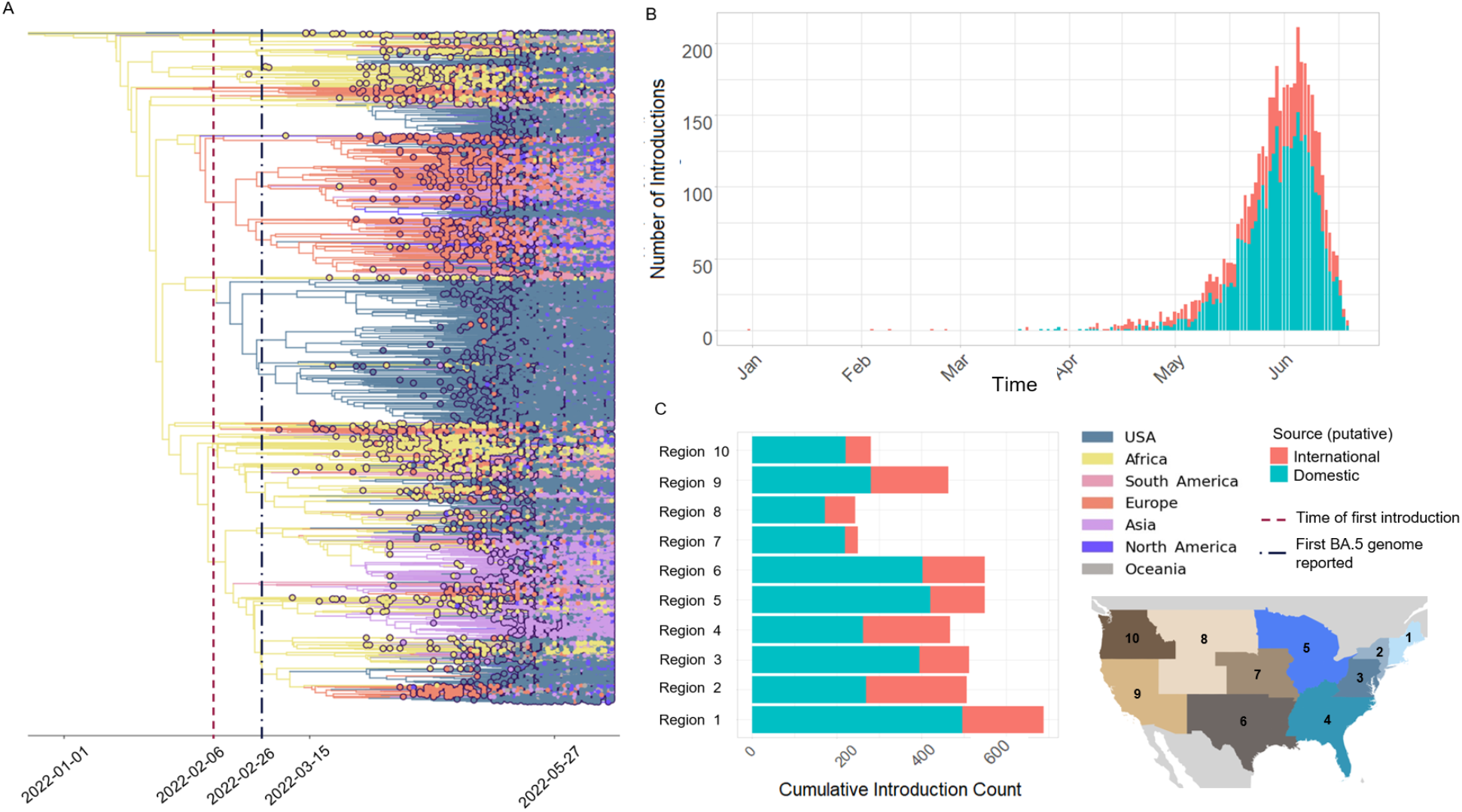
(February to June 2022) (A) Result of phylogeographic analysis of BA.5 emergence using 18,529 U.S. and global sequences. Dotted lines indicate the date of the inferred first introduction and the first sample of BA.5 in the U.S. (B) Comparison of the timeline of introductions by foreign vs domestic origins, with Orange as foreign introductions into U.S. regions and Blue for introductions between U.S. domestic regions, shaded area indicates early emergence period prior to May 15, 2022. (C) Total introductions into the 10 HHS regions of the U.S. during BA.5’s emergence period, split by their domestic or international origin.

**Figure 3:**
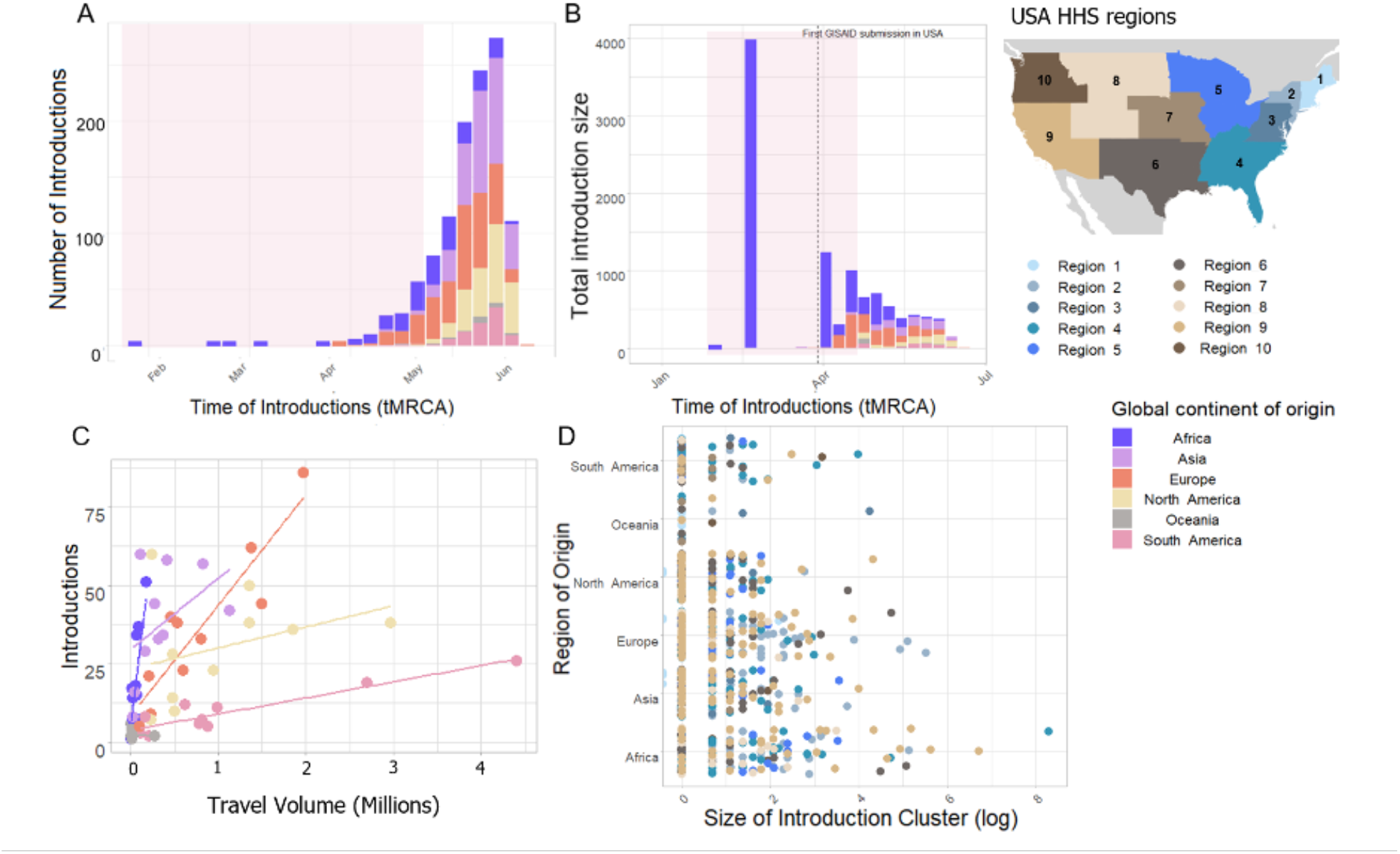
Spatiotemporal dynamics of BA.5 lineage foreign introductions to the U.S (February to June 2022). (A) Timeline of BA.5 foreign introduction events to the U.S., by continent, shaded area indicates early emergence period prior to May 15, 2022. (B) Total introduction cluster size of BA.5 foreign introduction events into the U.S. during variant emergence period, the shaded area indicates early emergence period prior to May 15, 2022 (C) Number of introductions into the ten HHS regions of the U.S. versus international travel volume, by continent, with x-axis representing numeric travel volume into the U.S. from a global region, and y-axis the number of introductions from the global region into the U.S. Each dot represents travel from a continent into a specific HHS region. Lines are regressions of travel volume against introduction number for each continent of origin. (D) Introduction cluster size into the 10 HHS regions of the U.S., by global continent.

**Figure 4:**
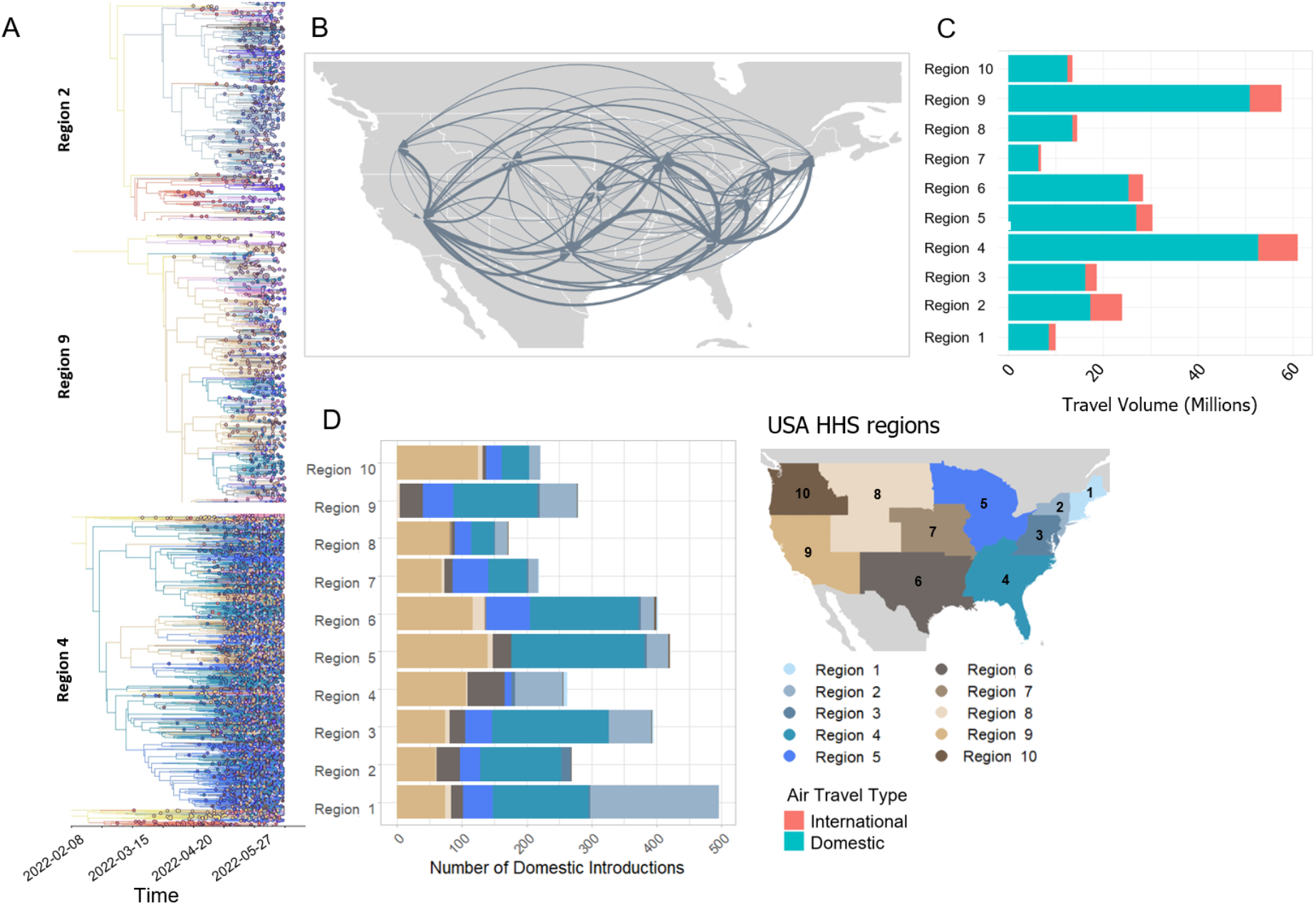
Spatiotemporal dynamics of BA.5 domestic introductions between the U.S. HHS regions in February to June 2022 (A) Time-scaled phylogenies of major U.S. clades, colored by the 10 U.S. HHS regions and labeled by their rooting (B) Map showing viral movements between regions within the U.S. with thickness of the lines indicates the prevalence of the movement across the maximum clade credibility tree and arrows displaying directionality of introduction. (C) Adjusted domestic and international air travel volume into the 10 U.S. regions (D) Number of domestic introductions into each region, colored by their origin within the U.S.

We observed a change over time in the relative dominance of continents as origins of BA.5 introductions into the U.S. (**Figure 3A, 3B**). Introductions from Africa, despite only representing 14.7% of total BA.5 international introductions, comprised 41.9% of all international introductions before mid-May 2022. These introductions were at a high rate into all 10 U.S. regions, despite very low travel volume into the U.S. from countries in Africa (**Figure 4C, 3C**). Indeed, Africa had the highest ratio of BA.5 introductions per travel volume, at approximately 0.3 introductions per 1000 unit of incoming air volume (**Figure 3C**), likely because BA.5 originated there. As BA.5 prevalence rose globally, introductions from Europe and Asia became more important (**Figure 2A, 2B, 3A, 3B**), matching high travel volume. Therefore, early emergence was determined by the geographical origin of the variant, with later introductions more connected to travel volume.

Finally, we examined the impact of timing on the size of international introductions. Between the first introduction of BA.5 into the U.S. in early February 2022 and its detection approximately three weeks later, there were five total introductions (**Figure 2A, 3A, 3B**). While four were singletons, one notable introduction in late February from Africa contained 3980 sequences – the largest during the entire emergence period in this dataset (**Figure 3B, 3D**). Cluster size was highest during early introductions and decreased over time during the BA.5 emergence period (**Figure 3B**). Notably, African introduction events, mostly earlier in the emergence period, tended to result in higher outbreak clade sizes, with nine clusters of over 100 sequences (**Figure 3D**). European introductions resulted in only four clusters of over 100 sequences, and no other global regions had clusters of such size (**Figure 3D**).

We therefore found two main phases to the emergence of BA.5 in the U.S.. Large introductions from Africa dominated the early emergence phase prior to May 2022. As prevalence increased globally, international introductions became more tied to air travel volume, hence there were more introductions from Europe and Asia. Due to a decrease in the susceptible population, and possible behavior change following an uptick in cases, European and Asian introductions were not able to grow as large as the earlier African events.

### Domestic movement of BA.5 in the U.S. was facilitated by key hotspots and population size

To understand BA.5 transmission within the U.S., we performed a discrete phylogeographic analysis, using the 10 regions of the U.S. defined by the HHS (**Figure 1C**). In total, we inferred 3137 within-country introductions across a single posterior tree, approximately 70% of total introductions across the whole period. Early international introduction was followed by substantial domestic transmission (**Figure 2A, 2B**) and all 10 HHS regions received more than 50% of their introductions from domestic sources (**Figure 2C**). These domestic movements grew in proportion throughout the emergence period and overtook the proportion of international introductions (**Figure 2B, 3B**), aligning with the high proportion (80%) of air travel which was domestic (**Figure 4C**).

There was no noticeable geographical structure within the phylogeny, i.e. sequences from different locations were intermixed, implying frequent inter-region transmission during the emergence period (**Figure 2A**). Inspection of the three largest and earliest U.S. clades, titled by their rooting in region 2 (including New York), region 9 (including California), and region 4 (including Florida) **(Figure 4A**) showed that locations that were close together tended to have more movement between them: region 2 and region 4 clades primarily saw transmission to other East Coast regions (**Figure 4A, 4B**), and region 9 clades mostly exported to other West Coast and West-Central regions (**Figure 4A, 4B**). Nonetheless, the interaction between regions 4 and 9, and to a lesser extent regions 2 and 5 (including Illinois), indicated the presence of coast-to-coast spread as an important mechanism in BA.5 emergence.

There were several key hotspots for transmission. All HHS regions had considerably higher introduction counts from regions 4 and 9 (**Figure 4D**). Rates of interaction between regions 4 and 9 and other regions were the highest in the country, together forming 71.6% of total domestic movements of BA.5 (**Figure 4D**). Correspondingly, regions 4 and 9 also had the highest volumes of air travel, both international and domestic (**Figure 4C**). Region 1 (New England) had the highest number of incoming domestic introductions and nearly 70% of such events came from regions 4 and 9 (**Figure 4D**). It therefore appears that the strong transmission between regions 4 and 9 underpinned BA.5 emergence in the US. We also theorize that region 1 was the top recipient of domestic introduction events due to the higher rate of interstate travel between regions 1, 2, and 3, as well as incoming air travel from other regions (**Figure 4B, 4C**).

To explore possible underlying drivers of viral movement across the U.S., we performed a linear regression between pairs of locations using population sizes and whether they share a border as predictors. We found that the population size of the origin location was a significant predictor for the number of viral movements between a pair of locations (p < 0.0001), in comparison with the destination population and whether the two locations shared a land border (p>0.1).

Overall, our findings demonstrate that the domestic introduction of novel BA.5 in the U.S. was characterized by cross-country interactions between key hotspots, and larger origin populations were positively associated with a higher proportion of introductions between the two regions.

## Discussion

As SARS-CoV-2 continues to spread in the U.S. and around the world, it is important to understand how new variants will disseminate. In this paper, we found that BA.5 was first introduced into the U.S. primarily from its geographical origin in Africa and then spread domestically from large populations and key hotspots, which are common between variants.

We found that the earliest BA.5 introductions into the U.S. came from Africa, despite low rates of air travel, showing the importance of a variant’s geographical origin. They were also much larger than later introductions, which is a common thread among the different waves of SARS-CoV-2 across the globe, despite different demographic and intervention contexts (13). As prevalence rose globally in the latter half of the emergence period, there was a higher proportion of introductions from Europe and Asia, potentially corresponding to higher travel volume (see Results and (20)). Similar dynamics have occurred before, notably with Delta variant introductions into the UK (11). This combination of the earliest introductions being the most important and later introductions coming from many places make international travel restrictions challenging to implement - even aside from ethical concerns (21) - as the speed required to prevent the most important early introductions from the origin (if this is even known) is unachievable in most settings.

Domestic transmission played a significant role in BA.5 dissemination in the U.S. While rates of inter-region transmission exceeded those of global importation across the whole emergence period, most domestic viral movement took place in the latter phase. The result shows widespread secondary transmission across the U.S. following initial international introduction. It also corroborates past findings of SARS-CoV-2 transmission being driven by domestic dynamics (15,17). That domestic BA.5 spread within the U.S. was significantly associated with population size of the origin location fits with past descriptions of COVID-19 transmission starting from large urban centers into other areas (22,23), and, along with geographical proximity being somewhat important, fits a classical gravity model of disease transmission (24).

Cross-country interactions between region 9 (California and other southwestern states), region 4 (Florida and other southeastern states), and region 2 (New York and surrounding area), highlight the role of specific hotspots in promoting BA.5 emergence. These three regions received the most introductions from Africa and contained the three largest and earliest U.S. clades, thus playing an important role in receiving and disseminating early BA.5 introductions. This is similar to the dissemination of Alpha variant (17), where New York City received the most introductions from the origin of the variant, followed by California and Florida. Therefore, we may expect these regions to be important in future variant introductions. Further, we found that region 1 (New England) was the highest recipient of domestic introductions, likely because New England has high interaction rates with two of the key hotspots (regions 2 and 4). We suggest that the identity of these primary hotspots (Regions 2, 4, and 9) was due to their important urban centers (e.g. New York City, Los Angeles, and Atlanta). These findings fit with Tsui et al (2022)’s description of early viral lineage movements between larger cities, followed by spatial expansion into nearby areas.

This study has some limitations. First, our subsampling method was reflective of the broader inequality in genomic surveillance, around the world and within the U.S. (25,26). We attempted to minimize these biases through subsampling and categorization into broader continents and HHS regions. Our tree being rooted in Africa, despite European sequences in particular overwhelming the global dataset, suggests that our attempts to mitigate this international bias were somewhat successful. Our categorization into larger regions (within and outside of the U.S.) may have introduced residual confounding, as we would be unable to gauge inter-state introduction events. Further, our definition of the variant emergence phase was made based on the frequency growth curve to filter for early BA.5 sequences, which we deemed more important to our research purposes, and might not properly reflect the true emergence time for a novel variant. Finally, we did not test other factors which may be driving the international introduction of BA.5 into the U.S., such as distance through air networks or income level.

As a large-scale study of Omicron lineage establishment, our findings support the role of phylogenetics in SARS-CoV-2 surveillance and contribute a phylogeographic framework for studying the emergence of other infectious pathogens in the U.S. As countries have lifted pandemic restrictions and the general population has a mosaic of immunity, the epidemiological landscape presents opportunities for positive selection of novel SARS-CoV-2 variants. Understanding the different dynamics of introduction in U.S. regions is important for timely and cost-effective policymaking, particularly for health authorities. Our methods may extend beyond SARS-CoV-2 and form a framework for phylogeographic analysis of large datasets to discern the spatiotemporal spread of other novel pathogens.

## Methods

### Dataset generation

To reconstruct the dynamics of variant introduction into the U.S., we assembled a dataset of BA.5 whole genome sequences sampled in the country and around the world during the inferred emergence period, estimated between February and June 2022. First, we downloaded all sequences with complete location and collection date metadata from GISAID with the pango lineage designation BA.5 (27). We then used Nextclade to remove sequences with a low-quality control score and genome coverage of less than 70%. To attempt to mitigate sampling bias, we categorized global BA.5 data by continent. Within the U.S., the genomic surveillance policy is largely decided by the individual state, causing potential bias in data by region (25,26). To ameliorate this disparity, we divided the country into ten regions, according to the locations of the ten regional offices of the Department of Human and Human Services (HHS, **Figure 1**).

We defined emergence as the period between an initial variant introduction event to the U.S., or events, leading to its spread and achieving a minimum frequency of 27% compared to other lineages in the relevant location. As there is not yet a standardized rate for variant emergence, we set this threshold after examination of BA.5 frequency over time in every region, as well as the total number of genomes from GISAID. More specifically, the 27% minimum frequency ensured BA.5 had been established across all global continents and U.S. regions but would not result in an intractably large final dataset.

To account for possible selection bias from this heterogeneity, we subsampled by week proportional to the population of each region. Specifically, we used the proportion of the region (either global continent or U.S. region) and multiplied it with the total number of BA.5 genomes, to find one fixed number of genomes to be selected every week for said region. The final dataset selected for analysis consists of 18,529 sequences, 9,350 from the U.S. and 10,258 from non-U.S. countries. For the emergence period, the earliest sample was collected on February 25, 2022, whereas the latest sample was collected on June 19, 2022. Table 1 shows the final count of sequences for all the respective regions of analysis:

**Table 1:**
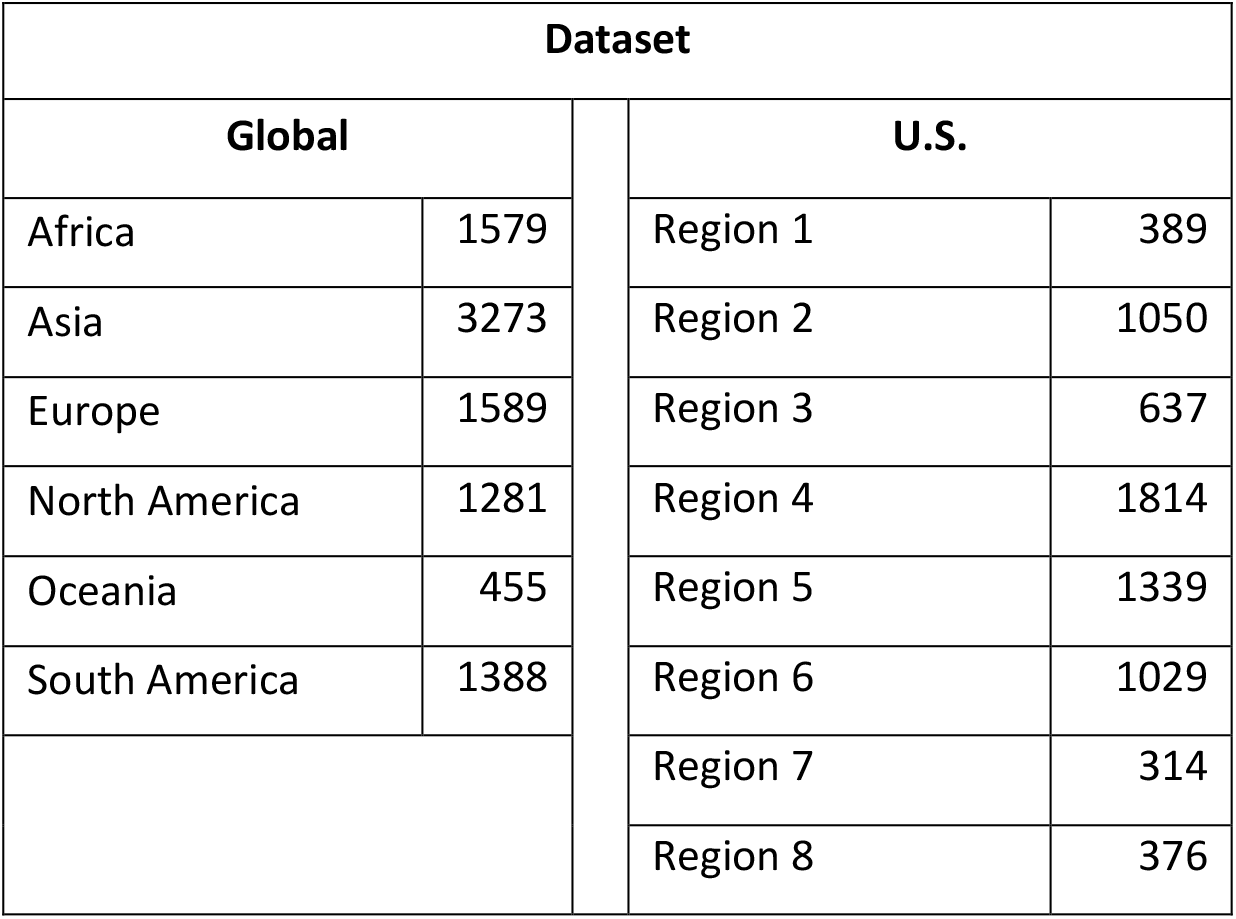
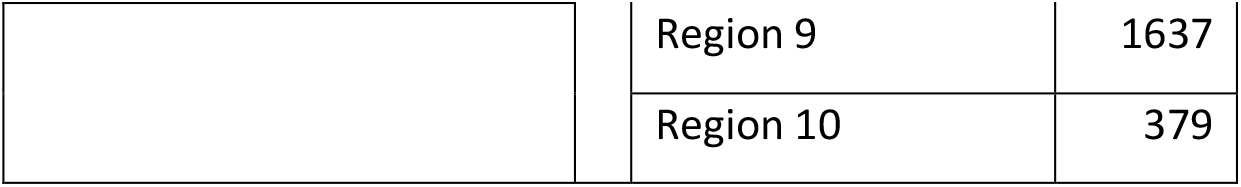
Final sequence dataset for BA.5 discrete phylogeographical analysis.

### Phylogeographic analysis

We performed a multiple sequence alignment using the Nextclade tool Nextalign (https://github.com/neherlab/nextalign), with Wuhan-Hu-1/2019 as the reference genome (GenBank accession: MN908947.3). Afterward, we constructed a maximum likelihood phylogenetic tree using IQ-Tree v2.2.2 (28) an HKY nucleotide substitution model (29) and outgroup rooting on MN908947. We assessed the temporal signal using TempEST v1.5.3 (30) and found that the dataset was from too short a timeframe to have a strong temporal signal. We were still able to prune molecular clock outliers using jclusterfunk v0.0.25 (https://github.com/snake-flu/jclusterfunk/) (31).

Given the large size of the genomic data set, we used an alternative tree likelihood function in BEAST v.1.10.4 which was developed for efficient estimation of large phylogenies (13,32). We used the maximum likelihood trees described above for the topological estimation, and approximately time-calibrated them using TreeTime v0.9.4 (33) to reduce the percentage of states needing to be discarded for burn-in.

Due to the low temporal signal in the dataset, we fixed the clock rate at 8×10E-4 substitutions/site/year, as in previous SARS-CoV-2 analyses (34–36). The non-parametric Skygrid coalescent model (37) was used with 23 grid points defined according to approximate equal intervals of the global emergence period shown above. We ran two Markov chain Monte Carlo (MCMC) chains for 1 billion iterations each, to ensure convergence to the same part of the posterior distribution. Tracer v1.7.1 was used to assess convergence upon run completion, with 10 percent of states discarded for burn-in (38).

Next, we picked one random tree from the post-burn-in posterior distribution of the previous analysis to use as the fixed tree in a discrete trait analysis. We performed analyses at two separate geographical scales - one at the global level using five continents - Africa, Asia, Europe, North America sans the U.S., Oceania, and South America - and the U.S. as a country, and one at the national level using ten HHS regions using the continental dataset as context. In both, we used an asymmetric continuous-time Markov chain (CTMC) to estimate transition rates between locations. Each MCMC ran for 2 million states and discarded 10% of the states for burn-in.

For the international analysis, we used custom Python scripts to estimate the average number of introductions across each tree in the posterior distribution and then selected a final tree that had the most similar number of introductions to that average number. For the domestic analysis, we chose a random tree in the posterior to maintain stability of within-US clades across the analysis. An introduction event is defined as the point in which a node is in a different location to its parent, either another continent into the U.S. for international introduction or between U.S. regions for domestic analysis. We do not allow for reintroduction within the same clade, so once the location changes to the U.S. said clade would be counted as one introduction event only. If a node in one subtree coincides with one from another subtree, we would only count the one with the older root and eliminate the other. The size of an introduction is the number of sequences that immediately follow a change in location within a node. We estimated the time of introduction as halfway between the first U.S./domestic location node and its parent. Visualizations were also generated using custom Python scripts, and trees were generated using the Baltic Python package (https://github.com/evogytis/baltic).

To examine some of the drivers of the domestic spread of BA.5, we constructed a linear regression model that incorporated geographic proximity and population. Within the model, the proportion of domestic introductions directionally between a pair of U.S. regions is the outcome, with two independent variables being the binary neighboring relationship between said pair and the numeric total population of the two regions. Population data was obtained from the U.S. Census Bureau.

### Travel data

To examine possible factors impacting BA.5 spread in different regions of the U.S., we collected monthly, international and domestic air travel data into U.S. states during our period of study (February - June, 2022) (39). The data are adjusted air passenger estimates. They are sampled data based on ticket sales, reporting from airline carriers, and assumed to represent 100% of the market. Adjusted travel volume represents the aggregate number of passenger journeys, not necessarily unique individuals. Passenger journeys are defined as the airline transport between original embarkment and disembarkment in the U.S. Both direct and indirect (i.e., connecting) flights are included.

## Data Availability

The flight travel volume data were provided by OAG Aviation Worldwide Ltd. OAG Traffic Analyser, Version 2.6.1 (http://analytics.oag.com/analyser-client/home; accessed 2023-04-24). These data were used under the United States Centers for Disease Control and Prevention (CDC) license for the current study and so are not publicly available. The authors are available to share the air passenger data upon reasonable request and with the permission of OAG Aviation Worldwide Ltd.

Genomic data was all obtained from GISAID, and a list of accession numbers can be found in **Table S1**.

## Acknowledgements

We thank Anne Hahn and Nicholas Chen for their help with this study. This project is supported by the Centers for Disease Control and Prevention (CDC) Broad Agency Announcement Contracts 75D30122C14697 and 75D30120C09570 (NDG). We thank everyone who has contributed genomic data to GISAID which makes work like this possible. To view the contributors of each individual sequence with details such as accession number, Virus name, Collection date, Originating Lab and Submitting Lab and the list of Authors, visit 10.55876/gis8.240620dg.

The findings and conclusions in this research are those of the authors and do not necessarily represent the official position of the Centers for Disease Control and Prevention.

