## Supplementary Information for "Large-scale genomic analysis of SARS-CoV-2 Omicron BA.5 emergence in the United States"

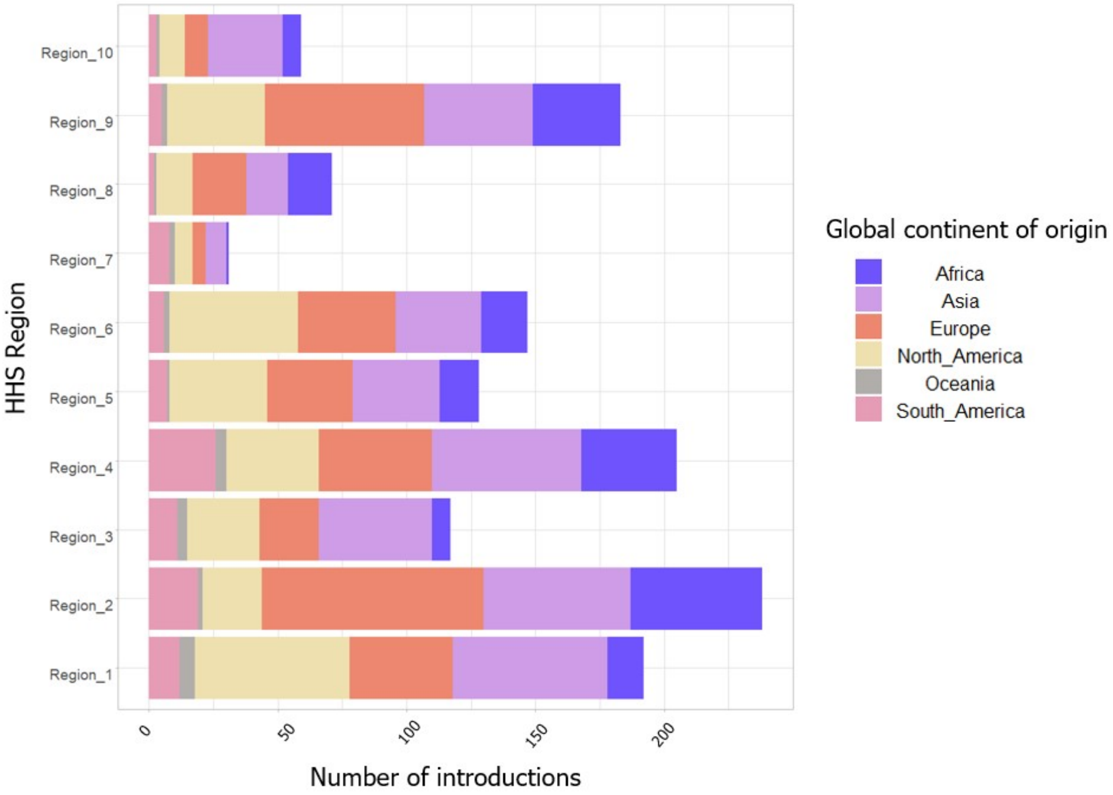


*Figure S1: Number of international introductions into the ten HHS regions of the U.S. during BA.5 emergence period, by global continent.*

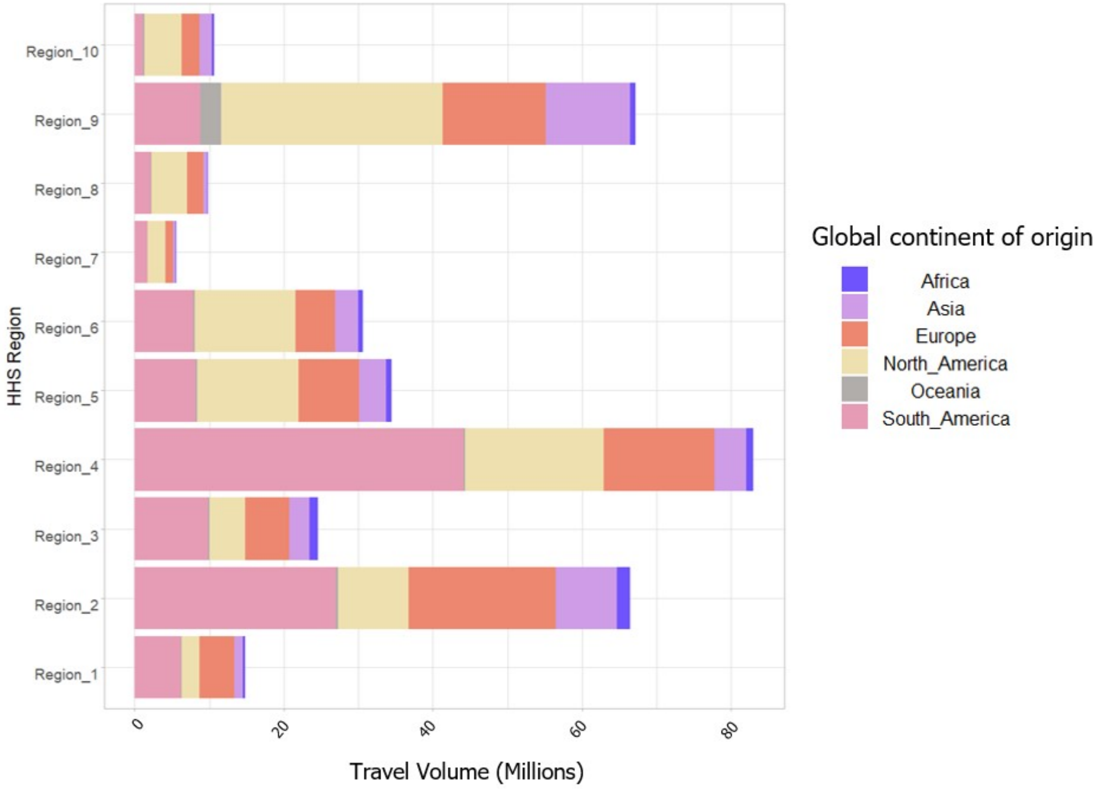


*Figure S2: Travel volume into the ten HHS regions of the U.S. during BA.5 emergence period, by global continent.*
